# Obstructions in Vascular Networks. Critical vs Non-critical Topological Sites for Blood Supply

**DOI:** 10.1101/018960

**Authors:** Aimee M. Torres Rojas, Alejandro Meza Romero, Ignacio Pagonabarraga, Rui D.M. Travasso, Eugenia Corvera Poiré

## Abstract

We relate vascular network structure to hemodynamics after vessel obstructions. We consider tree-like networks with a viscoelastic fluid with the rheological characteristics of blood. We analyze the network hemodynamic response, which is a function of the frequencies involved in the driving, and a measurement of the resistance to flow. This response function allows the study of the hemodynamics of the system, without the knowledge of a particular pressure gradient. We find analytical expressions for the network response, that explicitly show the roles played by the network structure, the degree of obstruction, and the geometrical place in which obstructions occur. Notably, we find that the sequence of resistances of the network without occlusions, strongly determines the tendencies that the response function has with the anatomical place where obstructions are located. We identify anatomical sites in a network that are critical for its overall capacity to supply blood to a tissue after obstructions. We demonstrate that relatively small obstructions in such critical sites are able to cause a much larger decrease on flow than larger obstructions placed in non-critical sites. Our results indicate that, to a large extent, the response of the network is determined locally. That is, it depends on the structure that the vasculature has around the place where occlusions are found. This result is manifest in a network that follows Murray’s law, which is in reasonable agreement with several mammalian vasculatures. For this one, occlusions in early generation vessels have a radically different effect than occlusions in late generation vessels occluding the same percentage of area available to flow. This locality implies that whenever there is a tissue irrigated by a tree-like in-vivo vasculature, our model is able to interpret how important obstructions are for the irrigation of such tissue.

## Introduction

Occlusion of tubes has always represented a problem. From engines and filters to arteries and bronchia, we can find countless systems where a reduction of the fluid flow in a particular site due to the presence of an obstacle, results in the partial or total failure of a process. Occlusion of bio-tubes in the human body represent an important issue in many diseases, and the circulatory vascular network is particularly vulnerable to obstructions. For instance, after an occlusion in the arteries, blood flow decreases, and, in critical cases, is effectively suppressed downstream. Such decrease of flow may have serious consequences at different levels, affecting oxygen and nutrient delivery to a tissue, or implying an increase in the stress over the heart muscle [1]. One dramatic example of the pathological effect of vascular obstructions is a retinal artery occlusion. Such an occlusion by a blood clot withdraws the nutrient and oxygen supply from the retinal cells and may render an individual blind from an eye within a few hours [2, 3]. In this scenario is of the utmost importance to identify specific sites in a vasculature where a partial obstruction can dramatically affect blood supply.

Various simulations of flow around partial obstructions in vessels exist in the literature [4–12]. Such analyses aim at describing in detail the flow patterns around occlusions, such as the velocity at different points inside a vessel, the existence of vortices, or the values of wall shear stress at different locations [5–7, 9–12]. Depending on the interest of each particular study, they may account for 3-dimensionality, elasticity or viscoelasticity of the vessels, inclusion of non-linear convective terms, and the effects on flow of bifurcations, to name a few.

The enormous amount of work involved in such computations is necessary when one wants to describe specific zones of a vasculature, and to answer detailed questions regarding flow profiles around obstacles, stenosis, bypasses, bifurcations, or flow in the aortic arch. These complex computations are able to predict how the waveforms of pressure and flow change in certain vessels due to obstructions, stenosis or vessel suppression at particular sites [4, 8, 13]. Sophisticated models are also very interesting from a theoretical and computational point of view. However, they involve too many variables to allow for the derivation of analytical expressions when one is interested in knowing the effect that obstructions have on the overall flow throughout an entire network. Analytical expressions might be very powerful and are potentially useful clinically, where a reduced number of parameters is often appreciated.

Knowledge about the structure of vascular networks, is key to predict the flow after alterations in the vasculature, e.g. after the growth or introduction of new vessels [14, 15] or after the partial occlusion of vessels in the system [8, 11, 12, 16]. The correspondence between local structural network information and global flow through a network after vascular alteration, was put forward in the work of Flores et al [14]; the simplicity of the model allowed for analytical expressions that in turn lead to conclusions not attainable otherwise. For instance, it was demonstrated that the increase of flow in the network after the growth of new vessels in the form of anastomosis, is determined by the morphology of the vasculature in a small neighborhood around the place where the new vessels are included. Other processes that regulate vessel width, such as the myogenic effect, were shown to have a very small qualitative effect in how the increase in response depends on the localization of the anastomoses.

The purpose of the present study is to relate the basic, generic characteristics of an arterial vasculature with the flow that goes through it after anatomical variations caused by obstructions or vessel suppression occur. We deliberately keep a reductionist approach in order to obtain analytical expressions for the system response in which the roles played by the network structure, the degree of obstruction, and the geometrical place where obstructions occur, can be clearly identified.

We study flow in three types of networks: one constituted by identical vessels, a second one in which radii are given by Murray’s law, and a third case in which large changes in resistance exist within the network. We show how the underlying network can lead to radically different behaviors of the hemodynamic response and identify structural features present in tree-like vasculatures that are critical for the overall capacity of the network to supply blood after obstructions. We demonstrate that our results are local in the sense that they depend on the network structure around the place where obstructions occur. This implies that whenever there is a tree-like network in an in-vivo vasculature, our model is able to interpret the effect that an obstruction has on flow.

## Background

Recently, a model has been introduced in order to study viscoelastic flow in a network of tubes [17]. This model consists of a tree-like network in which rigid vessels bifurcate always into identical vessels giving rise to identical branches of the network. At each bifurcation step, the possibility of changes in the cross sectional area and the length of the vessels is allowed. Each level (or generation) of the network is constituted by vessels with the same length and cross section. Segments belonging to the same level are labeled with the same index. Outer levels of the network are the ones that are closer to the main branch, inner levels are the ones that are the result of several successive bifurcations.

The model considers a linear viscoelastic fluid with the rheological characteristics of blood [18] in a range of shear rates where there is no shear thinning, and analyzes the network hemodynamic response to a time-dependent periodic pressure gradient. A Maxwell fluid [19] is used for this study, but the formalism can be easily generalized to consider any linear viscoelastic fluid [20]. By considering mass conservation, and assuming that the total pressure drop is the sum of individual pressure drops, the dynamic response of the network, *χ*(*ω*), is written in terms of the dynamic permeability of individual vessels *K*_*i*_(*ω*) as

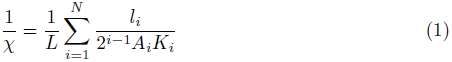

The sum is over the network levels, *A*_*i*_ and *l*_*i*_ are respectively the cross sectional area and the length of the vessels at the *i*-th level, L and N are the total length of the network and the total number of levels, respectively. The dynamic permeability for a vessel of radius *r*_*i*_ is 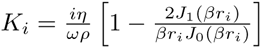 where *J*_0_ and *J*_1_ are Bessel functions of order zero and one, respectively, and 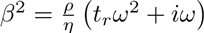, where *ρ*, *t*_*r*_, and *η* are the density, relaxation time and the fluid viscosity respectively. In order to apply Eq. (1) to a particular network of vessels, the network geometrical characteristics, namely, the number of levels-that determine the number of vessels-, lengths and radii, are required.

The network hemodynamic response relates viscoelastic flow and pressure drop in frequency domain [14, 17]. In order to have it explicitly in time domain one needs to specify a time dependent pressure gradient. As the equations are linear, we can obtain the fluid response to any time-dependent pressure gradient as a linear superposition of sinusoidal modes. For a single-mode time-dependent pressure drop Δ*p* = Δ*p*_0_ cos(*ω*_0_*t*), the volumetric flow as a function of time is given by

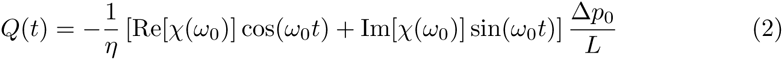

where the real and the imaginary parts of the response function *χ* (Eq. (1)) give the flow in-phase and out-of-phase with the pressure gradient, respectively [21]. Eq. (2) puts forward the importance of the dynamic response as a measurement of the resistance to flow. For systems driven at biologically relevant frequencies like the ones imposed by the heart, the imaginary part of the response is often negligible compared to its real part and the real part of the response gives one the proportionality factor between pressure gradient and flow in time domain. The response function allows one to study the hemodynamics of the system, without the requirement of considering a particular pressure gradient. Our results are presented at 1.5 Hz, which is the resting heart rate of the dog [22, 23]. At such low frequencies the network response (and total flow) is almost indistinguishable from the steady-state regime where the response is real. However, we keep the formalism as general as possible to make it applicable when external frequencies are imposed [21]. We use parameters for normal dog blood [24], *ρ* = 1050 kg/m^3^, *η* = 1.5 × 10^−2^ kg/(m s) and assume that the relaxation time is similar to the one reported for human blood: *t*_*r*_ = 1 × 10^−3^ s [18, 25].

## Model for obstructions in a tree-like network

The vascular system of mammals has a complex topology. However, there are several places in the body in which tree-like networks at different scales irrigate certain regions or tissues, from the tree-like networks resulting from successive bifurcations of large arteries that irrigate the limbs, to the tree-like networks characteristic of the microvasculature that irrigates the eyes.

We use an electrical analogy in which the resistance of each vessel is given by 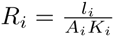. Medical and biological literature frequently report the fraction, *f*, of the total cross sectional area that has been obstructed. Accordingly, we consider the area of an obstructed vessel, 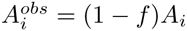. The radius of the obstructed vessel, 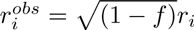, modifies its permeability and its corresponding resistance. We consider that obstructions occur in half of the branches of the same tree level as illustrated in Fig. 1A. Although such an obstruction pattern does not correspond generically to physiological conditions, it helps to highlight the impact of vessel geometry for equivalent obstructions, that is, those which block the same percentage of cross sectional area regardless of the level in which they occur.

**Figure 1.**
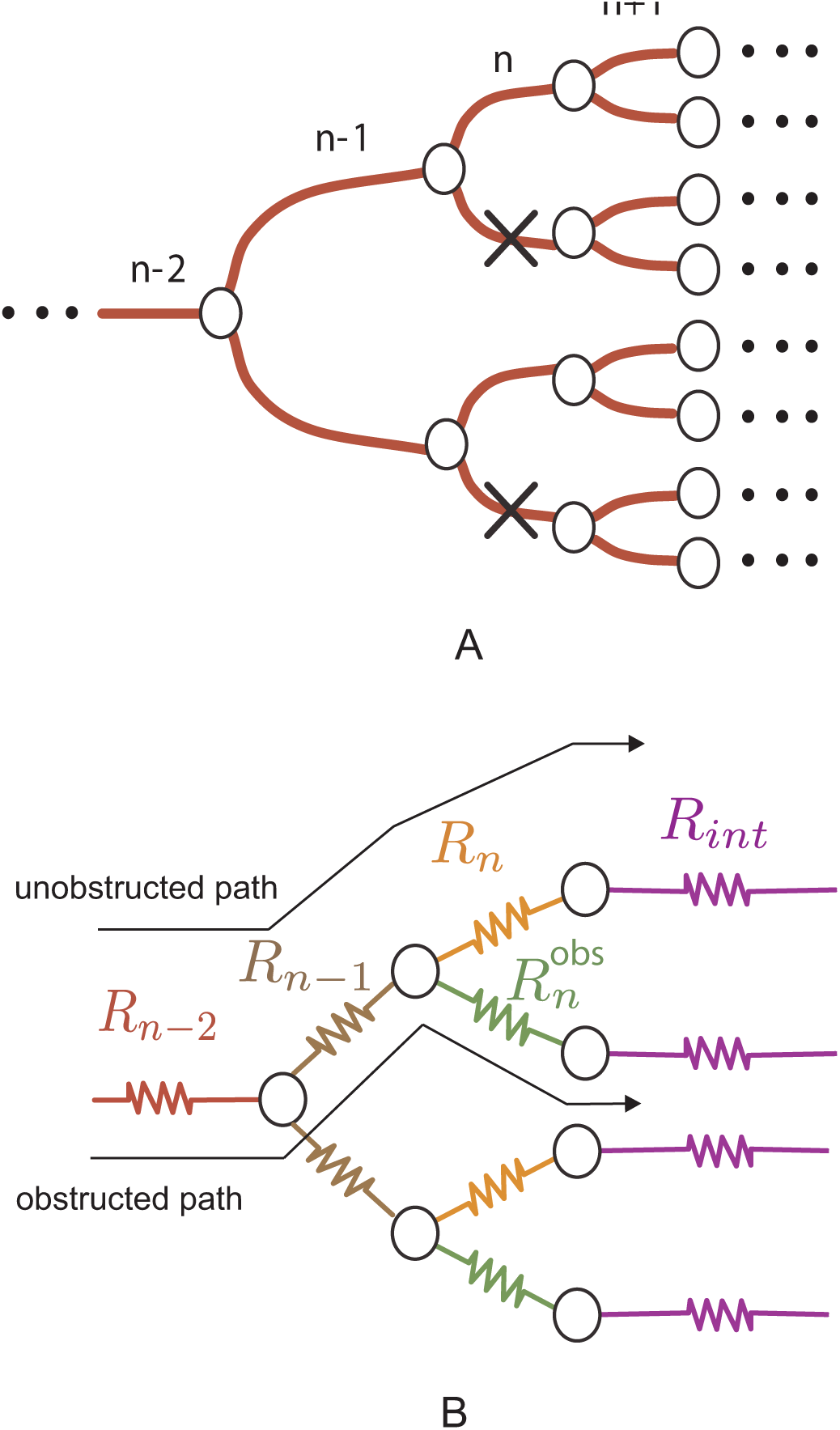
Model for obstructions in a tree-like network. A: Illustration of a network with obstructions at level *n*, indicated by crosses. B: Electrical analogy for a *N*-level network with occlusions at level *n*.

The total resistance for an *N*-level network obstructed at level *n* is given by

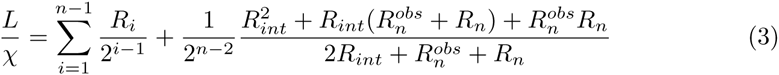

Where 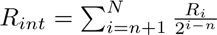 and 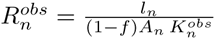 where 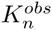 is the permeability of an obstructed vessel; Fig. 1B shows the electric analogy for a segment of a network with obstructions at level *n*.

Although we will focus on the overall behavior of the network, the analytical approach can predict the local flow at each of the network vessels. We will characterize the impact of vessel obstruction on tree-like networks by focusing on two different types of paths. We will consider *unobstructed paths*, those which cross the network without moving along any obstructed vessel, and *obstructed paths*, when an obstructed vessel is crossed at some point in the network.

## Obstructions in a network with equal vessels

We first treat the case of a network in which all vessels have approximately the same radius, which is the case of several networks at the arteriole level, and approximate it with a bifurcating network of equal vessels with resistance *R*_1_. We find that in this case, the effect caused by occlusions is relatively small when it happens in the inner vessels, and it is relatively large when it happens in the outer vessels. The network response increases monotonically with the level number *n* in which occlusions occur (see Fig. 2).

**Figure 2.**
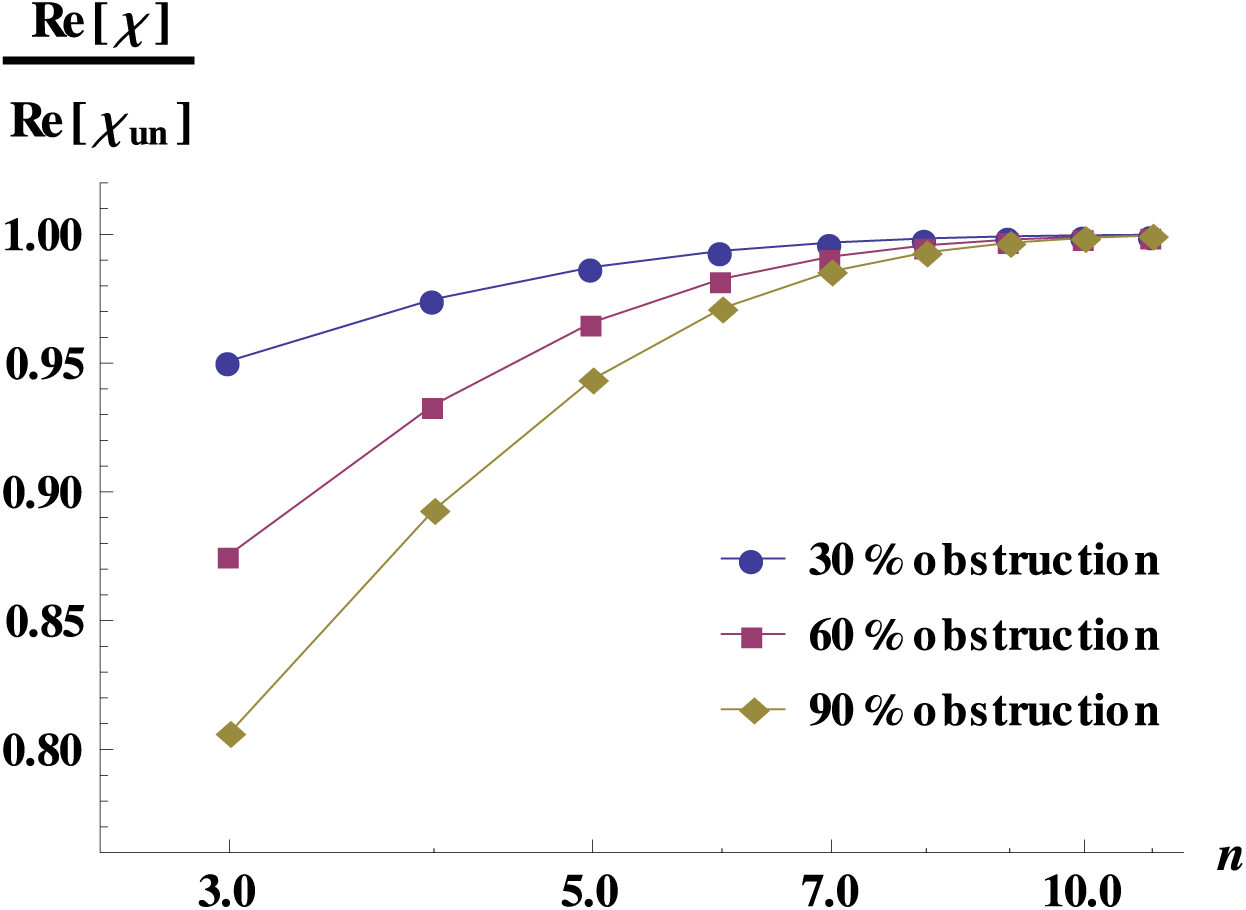
Dynamic response for obstructed networks with equal vessels. Dynamic response for an 11-level network as a function of the level *n* in which obstructions occur. It is important to note that each point in this figure corresponds to a different network since we obstruct only one level at a time. The normalization is done with the network response without occlusions. The effect of the obstruction is more dramatic in the outer levels of the network. In this calculation, the vessels have the typical dimension of the dog arterioles (*r* = 1 × 10^-^5 m and *l* = 2 × 10^-3^ m).

Physically, this implies that for a healthy tissue irrigated by a tree-like network, occlusions are more dangerous when they occur in vessels of early generations since blood supply is dramatically decreased, as illustrated in Fig. 3.

**Figure 3.**
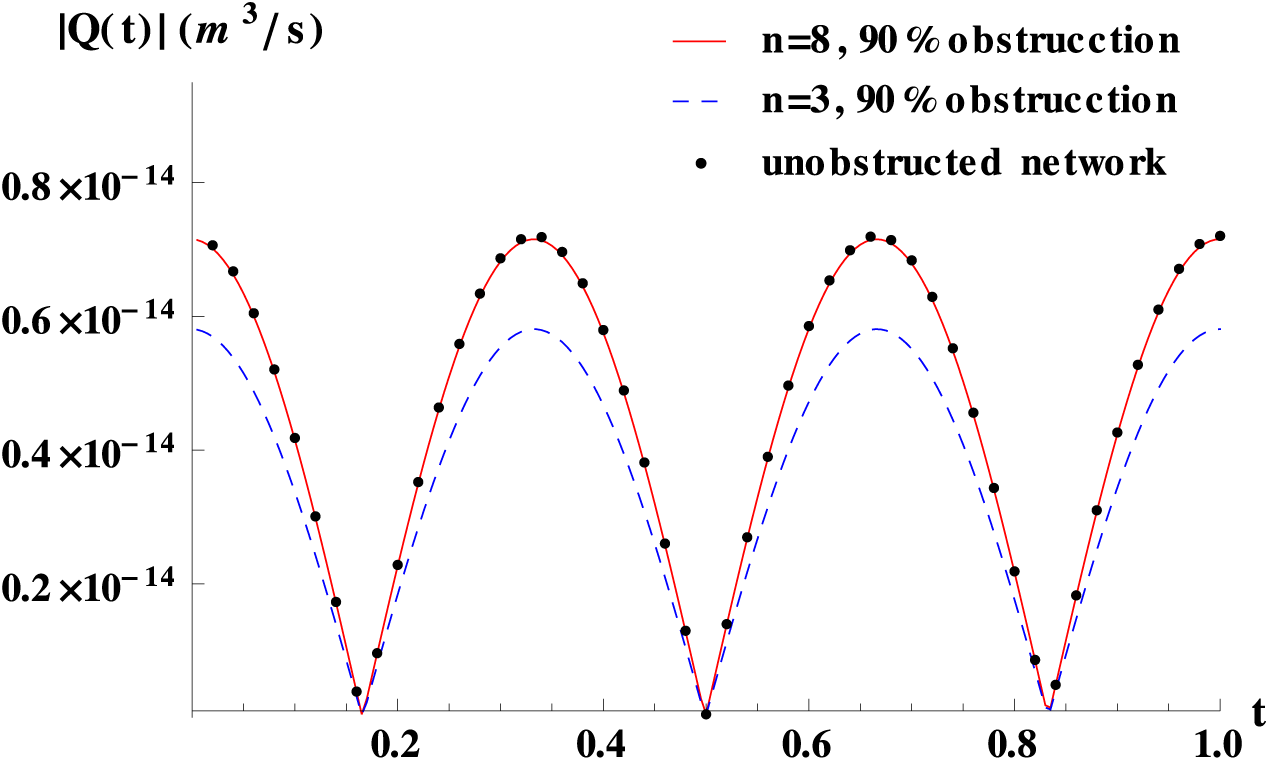
Time-dependent flow for obstructed networks with equal vessels. Blood flow for an 11-level network with obstructions of 90% at levels 3, 8 and with no obstruction (reference). The sharp decrease in flow after obstructions at the outer level is clear. The network used was the same as in Fig. 2. The total pressure drop was set to 110 Pa.

A mathematical analysis similar to the one presented in [14] for anastomosis, allows us to have an analytical approximated expression for the dynamic response of a network of equal vessels, *χ*, obstructed by a fraction of area *f* at level *n*,

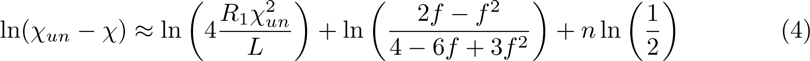

In this expression *χ*_*un*_ is the response of the unobstructed network. The last term in Eq. (4) is related to the anatomical place, *n*, where the obstructions occur and ln 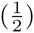 is due to the bifurcation nature of the tree. The constant terms on the right hand side of Eq. (4), are independent of *n* and depend only on the unobstructed network features and on the fraction of the occluded area *f*. Fig. 4 displays the remarkable good agreement between the numerical exact results and the analytical approximation for the dynamic response of a 20-level network, regardless of the level where the obstruction takes place.

**Figure 4.**
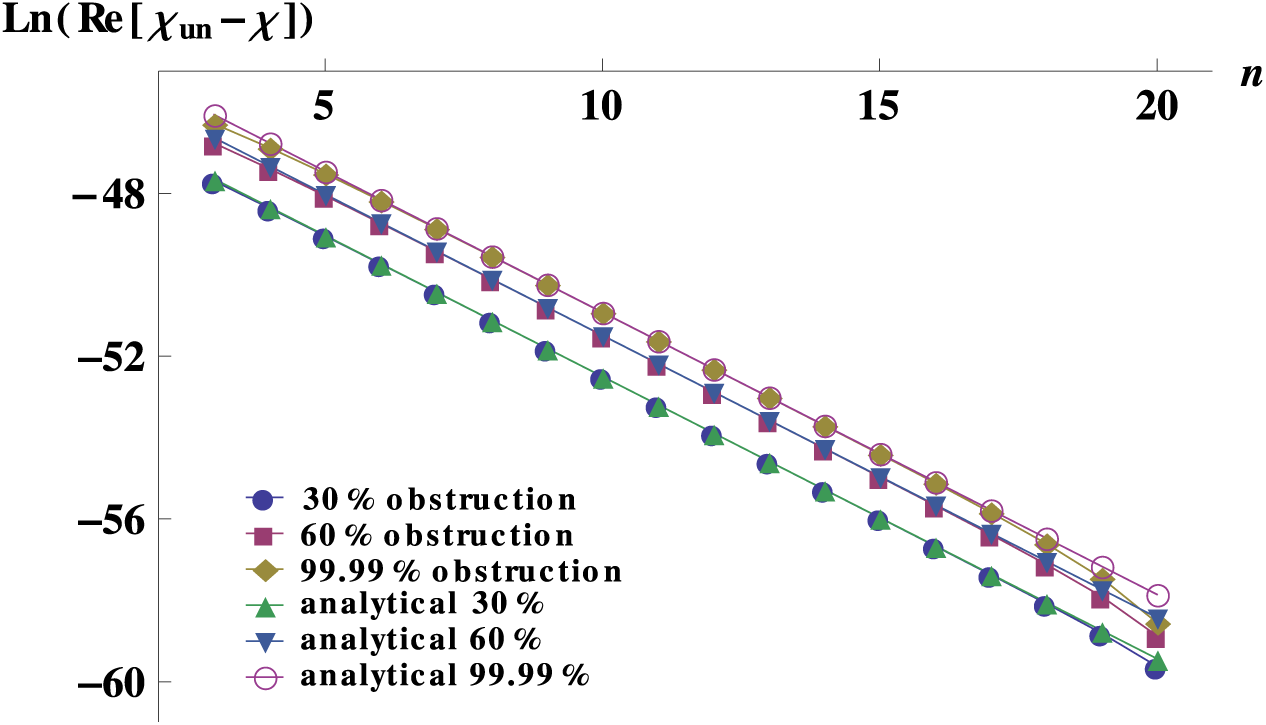
Analytical approximation and numerical solution of the response for obstructed networks with equal vessels. Analytical approximation and numerical solution of the quantity ln(Re[*χ*_*un*_ *- χ*]) for 20-level networks obstructed at level *n* as described in the text. The vessels have the typical dimension of the dog arterioles (*r* = 1 × 10^-^5 m and *l* = 2 × 10^-^3 m).

The theoretical prediction provides insight in the impact that the degree of obstruction and its location inside the network has in its global response; in particular the expression derived clearly shows that the change in the network’s response due to the presence of obstructions is highly determined by the structure of the unobstructed network.

It is very important to keep in mind that a global decrease of the total flow in a network, does not imply that all vessels have a smaller flow than in the absence of obstructions. For instance, Fig. 5 shows the local flow through the unobstructed and obstructed paths when obstructions that occlude 60% of the vessel section are placed in half of the vessels at level *n* = 3 of an 11-level network. The figure quantifies the relative increase (decrease) of the local flow through the unobstructed (obstructed) path. Because of flow conservation, the slopes in the log-linear plot in Fig. 5 for the flow through obstructed and unobstructed paths are identical [26]. Flow in the outer levels of the obstructed network is smaller than the reference curve, because the total flow (equal to flow at level 1) is smaller for an obstructed network than for an unobstructed one.

**Figure 5.**
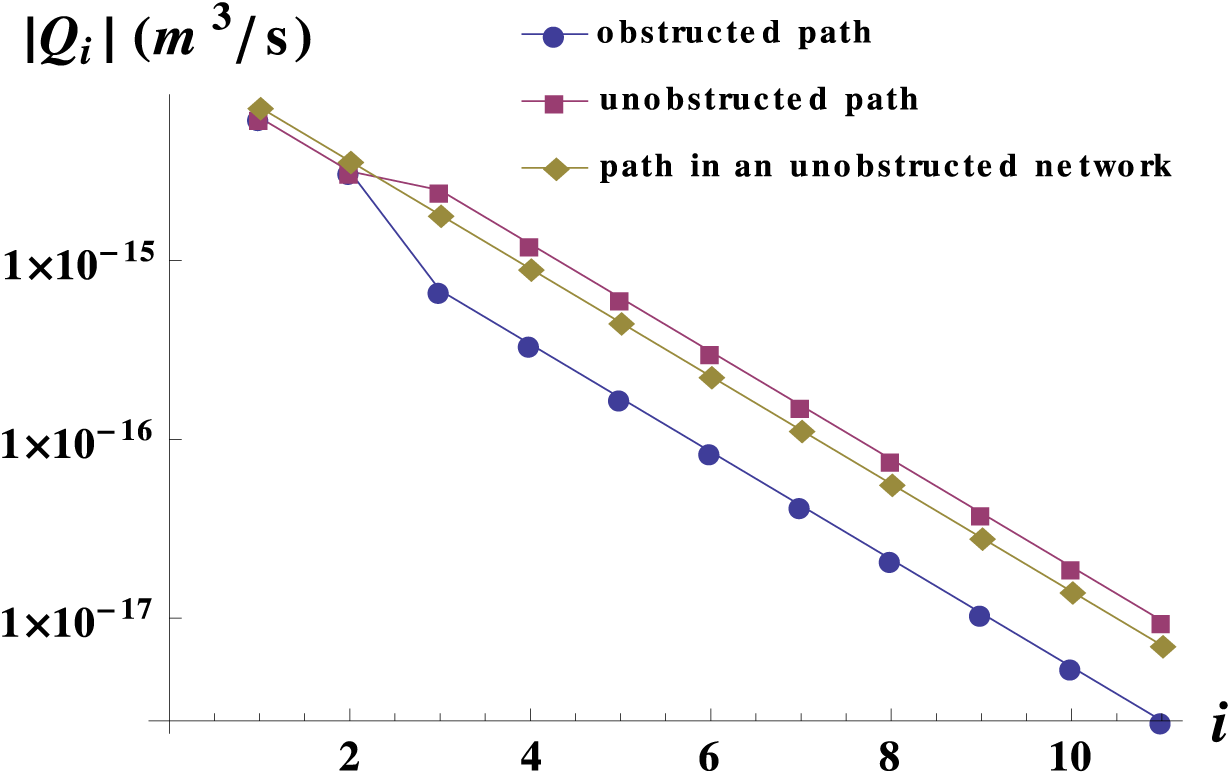
Flow in single vessels of an obstructed network with equal vessels. Flow in single vessels in logarithmic scale as a function of the level they belong to for a network constituted by 11 levels and with obstructions of 60% in area at level 3. The curves shown are: the flow in the unobstructed path, the flow in the obstructed path and a reference curve for the flow in a path of an unobstructed network. For language clarification see Fig. 1. Even though the total flow decreases with the obstructions, the flow in the non-obstructed vessels increases. The network used was the same as in Fig.2. The pressure drop was set to 110 Pa.

## Obstructions in a network with vessel radii that follow Murray’s law

Real vascular networks are composed of vessels of different radii and lengths; accordingly, they are characterized by changes in resistance from one level to the next one. For animal tree-like vasculatures, this normally implies an increase in resistance from one bifurcation level to the next one, because inner levels have smaller radii.

Assuming that the vascular system evolved to minimize the power required to maintain and circulate blood [27], Murray derived, in 1926, the relationship known as Murray’s law. This one relates the parent radius, *r*_*p*_, and the two daughters vessels radii, *r*_*d*1_,*r*_*d*2_, before and after a bifurcation, as

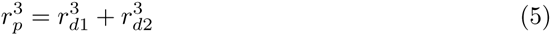

According to an extensive study on the validity of Murray’s law [28] and a review on vascular flow of reference [29], physiological studies showed that, barring some anomalies, a large part of the mammalian vasculatures have reasonable agreement with Eq. (5). According to [29], there is also a considerable mass of literature comparing physiological studies in animals other than mammals, and even in plants [30–33], that show good agreement with Murray’s law.

We therefore consider Murray’s law as an example of physiological relevance, in which our analytical results illustrate how to explain the different tendencies in the dynamic response in different sections of the network.

For our studies, we consider symmetrical branching, so in this case, radii of subsequent levels are given by

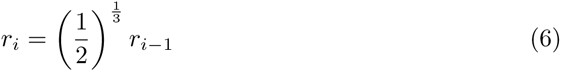

For lengths, we consider a power law decay with parameters that match the actual length of the aorta and the length of the capillaries of the dog circulatory system.

Figs. 6B left and 6B right, show the real and imaginary parts of the ratio of two sequential resistances of the underlying network, 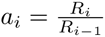 where 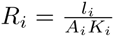The complex character of *a*_*i*_ is due to the fact that we are not working in a steady state, but at the frequency of the heart rate of the dog at rest. As for any realistic vasculature, the value of *a* changes along the network according to the lengths and radii of the vessels that compose it. A decrease in radii between subsequent levels produces an increase in resistance, on the other hand a decrease in length between subsequent levels, produces a decrease in resistance. It is therefore the interplay between this two quantities which will determine the value of *a*. It turns out that for a bifurcating network, the response will be qualitatively different whenever *a* is smaller or larger than 2, as we will see below.

**Figure 6.**
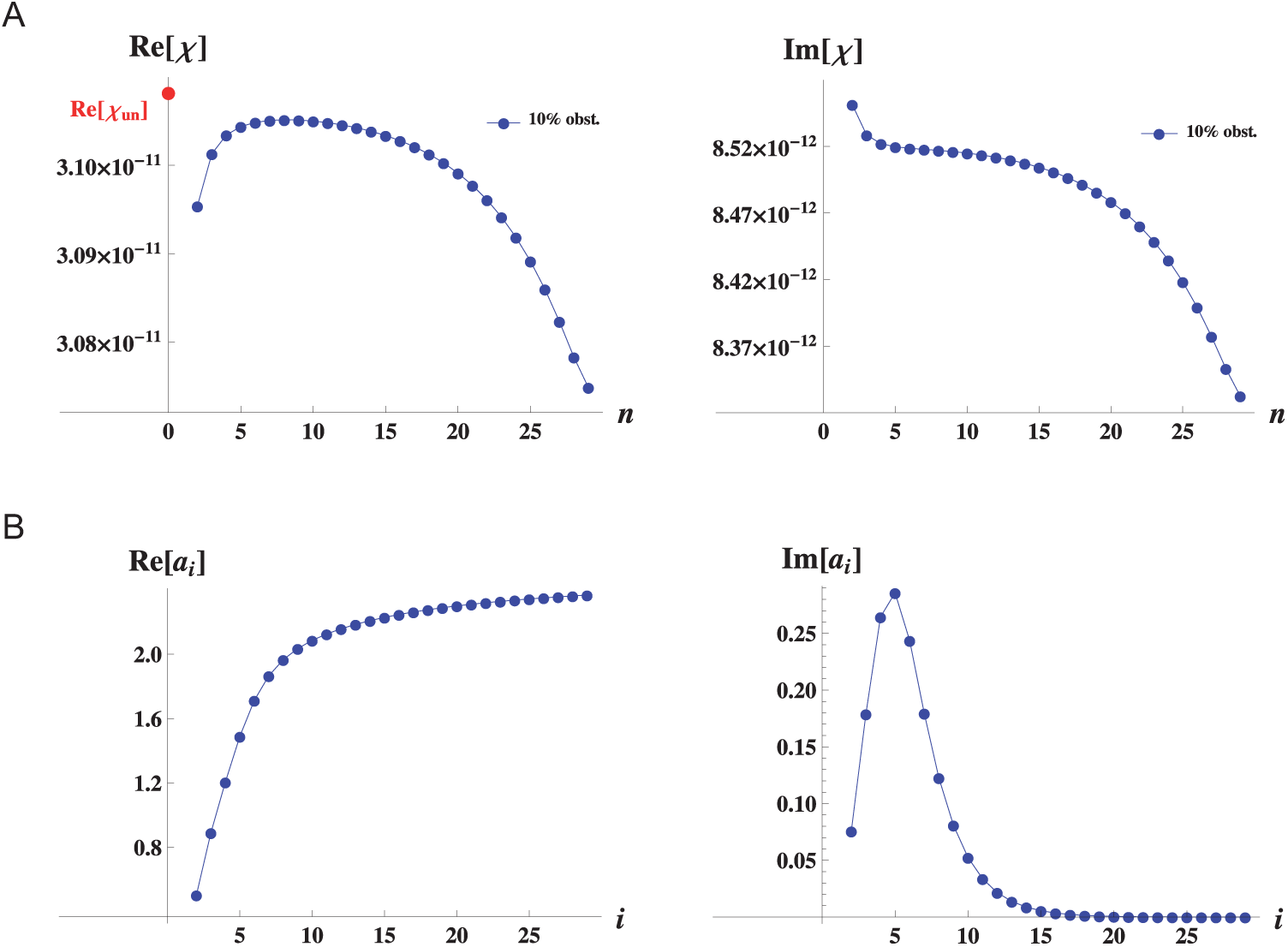
Dynamic response for a network with vessel radii that follow Murray’s law. A: Real and imaginary parts of the response of the network (in *m*^4^) as a function of the level *n* at which obstructions occur. B: Real and imaginary parts of the ratio of two sequential resistances 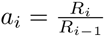 as a function of the level *i* of the underlying network. Note that we use the subindex *i*, whenever we refer to a property of the underlying network, we use the subindex *n* whenever we refer to the response of the whole network when obstructions occur at level *n*.

Figs. 6A left and 6A right, show the real and imaginary parts of the response as a function of the level *n* in which obstructions occur. As we can see from Fig. 6A left for large arteries, the real part of the response as a function of the level *n* in which obstructions occur, increases with increasing *n*. On the other hand, for the section of smaller vessels, the real part of the response as a function of the level *n* in which obstructions occur decreases with increasing *n*.

In order to gain insight into these results, we present analytical approximations for networks in which the ratio of subsequent resistances is less than two or larger than two. These ones agree well with numerical results whenever the real part of *a* is considerably larger than its imaginary part. They are given by:

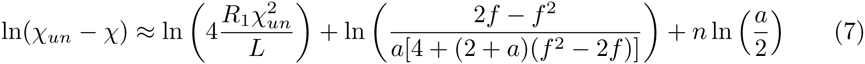

for *a <* 2, which reduces to Eq. (4) when *a* = 1, and

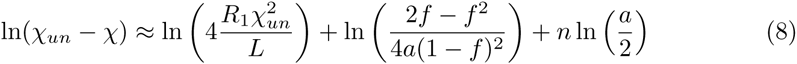

for *a <* 2.

The strongest influence of the underlying network without obstructions, on the network response when obstructions are present, comes from the term *n* ln 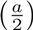 which gives a qualitatively different trend for *a <* 2 and for *a >* 2. In the first case *a <* 2, the response increases with increasing *n*. On the other hand, when *a >* 2, the response decreases with increasing *n*.

As we can see from Fig. 6A left for the outer levels, the response increases with increasing *n*, and as Fig. 6B left shows *a <* 2 on the same range. Likewise, for the inner vessels, the response decreases with increasing *n* and *a >* 2 on the same range.

For values of *a <* 2, just as in the case of equal vessels (that has *a* = 1) presented in the previous section, occlusions are more dangerous when they occur in vessels of early generations since blood supply is dramatically decreased. On the other hand, when *a >* 2, occlusions are more dangerous when they occur in vessels of late generations since blood supply is smaller than for instance at vessels around the middle of the network.

For the network presented here, that follows Murray’s law for radii and a power law for vessel lengths, two radically different behaviors are observed, one for external levels, and one for inner vessels. Our analytical results help us to build an intuition for the interpretation of the different behaviors of the response function in occluded vascular networks. Such behaviors are strongly dependent on the structure of the underlying network.

## Obstructions in a network with a jump in resistance

Finally, we consider a network for which vessels have a sudden jump in resistance. This corresponds to physiological conditions in cases where vessels of small radii branch from vessels of large radii. In particular, we consider a network for which we have a resistance *R*_1_ for *i ≤ k* and a resistance *R*_2_ for *i > k*, where *k* is a level close to the middle of the network. Therefore, the network has a jump in resistance 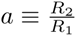 between levels *k* and *k* + 1. We obtain analytical approximations for real *a* that could be useful when one analyzes jumps in resistance in the arterial tree of mammals. The first case holds in the limit when 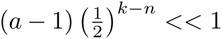 for *n* ≤ *k* and for a given *a* it is better the farther away from the jump obstructions are. In this case, we find that the network response is given by

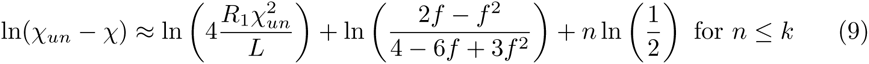

and

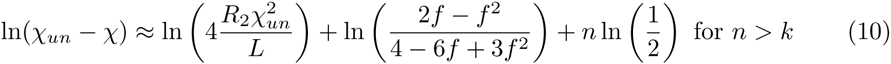

The right hand side of these expressions clearly highlights that the geometry of the underlying network, the fraction of obstructed vessels, and the geometrical place, *n*, where the obstruction is located contribute additively to the change in the network response. As examples of physiological relevance for which these analytical expressions could be useful, we find typical resistance jumps of the dog circulatory system [14] between main arterial branches and terminal branches, and between arterioles and capillaries. For the former example, the analytical approximation is good for degrees of obstruction up to 90%. For the later example, the analytical approximation is reasonably good for obstruction degrees up to 40%, Fig. 7A shows the exact and approximated values for the difference of the response for an obstruction degree of 40%.

**Figure 7.**
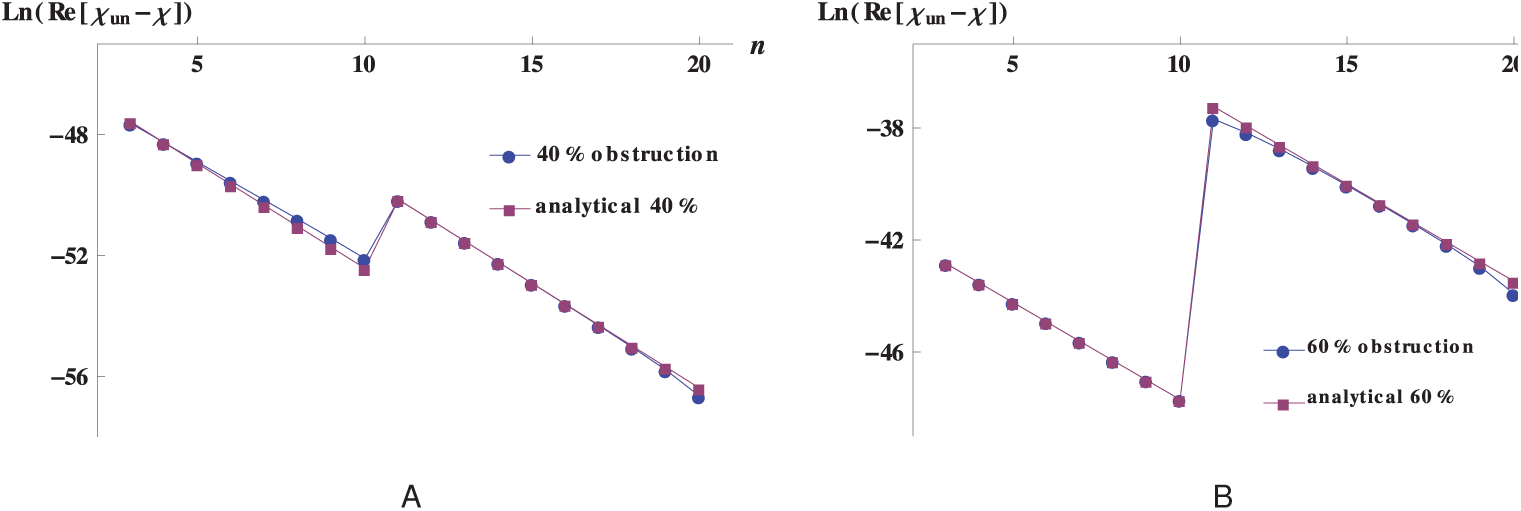
Analytical approximation and numerical solution of the response for obstructed networks with a resistance’s jump. Analytical approximation and numerical solution of the quantity ln(Re[*χ*_*un*_ *- χ*]) for 20-level networks with a jump in resistance between levels 10 - 11 obstructed at level *n* as described in the text. A: The vessels have the typical dimensions of the dog arterioles for *n ≤ k* and of capillaries for *n > k*. B: The vessels have the typical dimensions of the dog terminal branches for *n ≤ k* and of arterioles for *n > k*.

We have generalized these results for the case when a vasculature has several jumps, the approximated analytical expression is given by

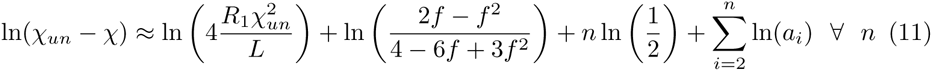

Here, *n* represents the level in which obstructions occur, and *a*_*i*_ is the ratio of two sequential resistances, 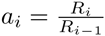. It is clear that whenever *a*_*i*_ = 1 there is no contribution to the sum in the last term of Eq. (11), and that for a network in which there are no jumps in resistance, Eq. (11) reduces to Eq. (4) with *a* = 1. This expression holds for jumps with small values of *a*_*i*_. The non-locality in this expression, given by the sum in the last term, does not prevent us from the possibility of interpreting locally the effect of obstructions. That is, in a local network, we may not know which is the precise value of the response-determined by jumps in resistances more external to the place where obstructions are present-, but relative to this value, we can quantify (through the term *n* ln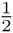) how, when there are no jumps in resistance, the response changes with the level *n* in which obstructions are. Also, we can identify (through the terms ln *a*_*i*_) the change in the response when obstructions are found in regions where the resistance has jumps.

We have worked out analytical expressions in the limit of larger resistance jumps, to be precise in the limit where, 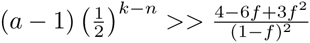. The network response can then be expressed as

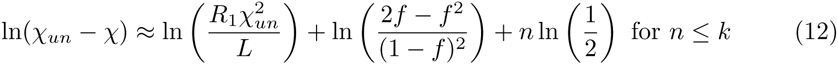

and

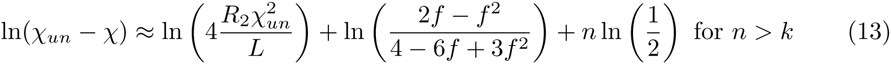

These expressions again show that the geometry of the unobstructed network, the fraction of vessel occluded area and the geometrical location of the obstruction in the network contribute additively to the change in the network response. As an example of physiological relevance, we show that these analytical expressions are useful for typical resistance jumps of the dog circulatory system [14] between terminal branches and arterioles for obstructions up to 60% as can be appreciated in Fig. 7B. The comparison between the numerical results and the theoretical approximation show again a very good agreement; for a given ratio in the jump in resistance, *a*, the analytic approximation is better the closer to the jump the obstructions are. Although it is possible to generalize the analytic expression for the network response for a series of jumps for which *a* is large, we have found no example in which this might be physiologically relevant.

The theoretical predictions for the jump between terminal branches and arterioles of Fig. 7B, are shown in a linear scale in Fig. 8, normalized by the response of the corresponding unobstructed network, for different degrees of obstruction, in order to better appreciate differences in response. The results show that obstructions occurring just after the jump in resistance diminish dramatically the network flow. At both sides of the geometrical place of the jump, we observe that the outer the level in which the obstruction takes place, the smaller the network response. We have checked that these features are generically true for any degree of obstruction. In order to illustrate how such variations in response, have a direct impact on flow we compare the network flow for obstructions that block 90% of the area at level 3 and obstructions that block only 45% of the area at level 11, as displayed in Fig. 9 where we also plot the flow for an unobstructed network as a reference. The figure clearly illustrates that relatively small obstructions in critical topological sites cause a much larger decrease on flow than larger obstructions in non-critical sites.

**Figure 8.**
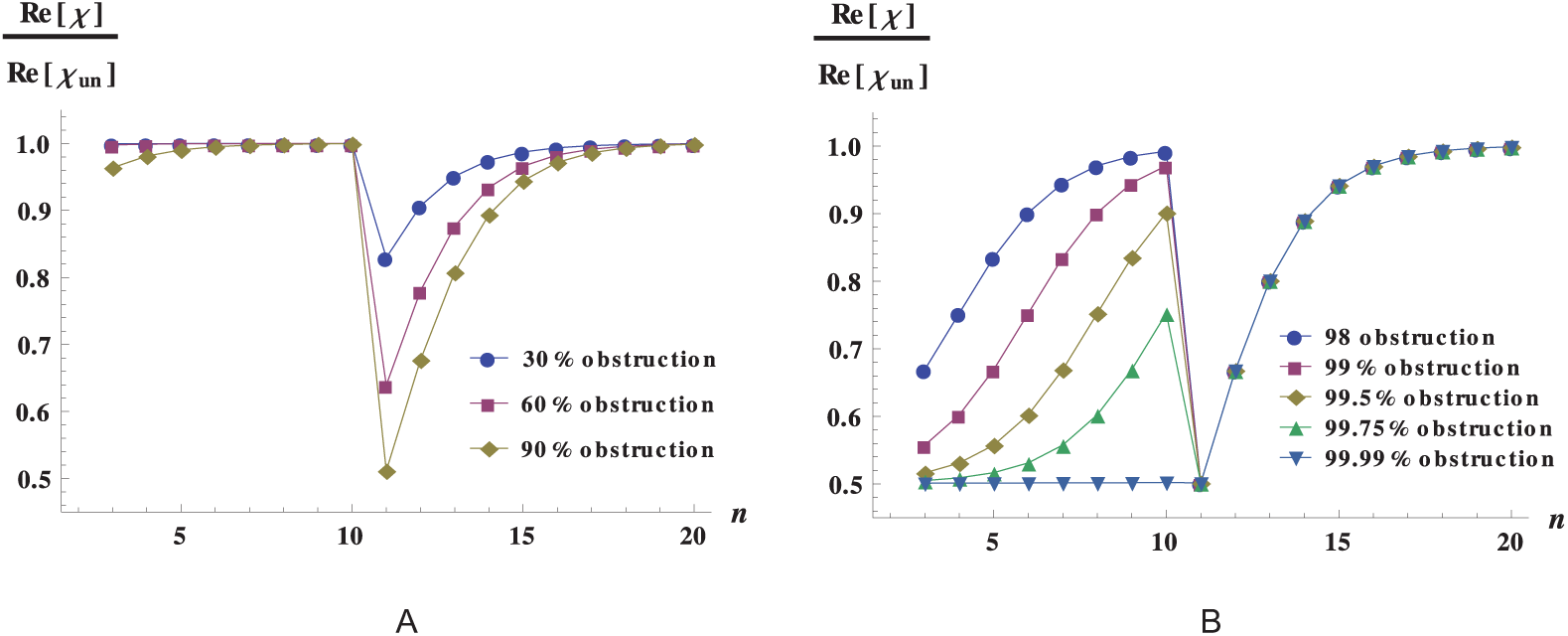
Dynamic response for the obstructed networks with a jump in resistance. Network response as a function of the level *n* in which occlusions occur. In this calculation we used a network with 20 levels, with an alteration in vessel radii and length between level 10 and level 11. We observe that obstructions at specific sites of network may result in a dramatic reduction of the response. The vessels of levels 1 to 10 have the typical dimension of the dog terminal branches (*r* = 3 × 10^-4^ m and *l* = 1 × 10^-2^ m) and the vessels of levels 11 to 20 have the typical dimension of the dog arterioles (*r* = 1 × 10^-5^ m and *l* = 2 × 10^-3^ m). A: Behavior dominated by the levels after the jump in resistance. B: Behavior for very large obstructions, dominated by the levels before the jump in resistance.

**Figure 9.**
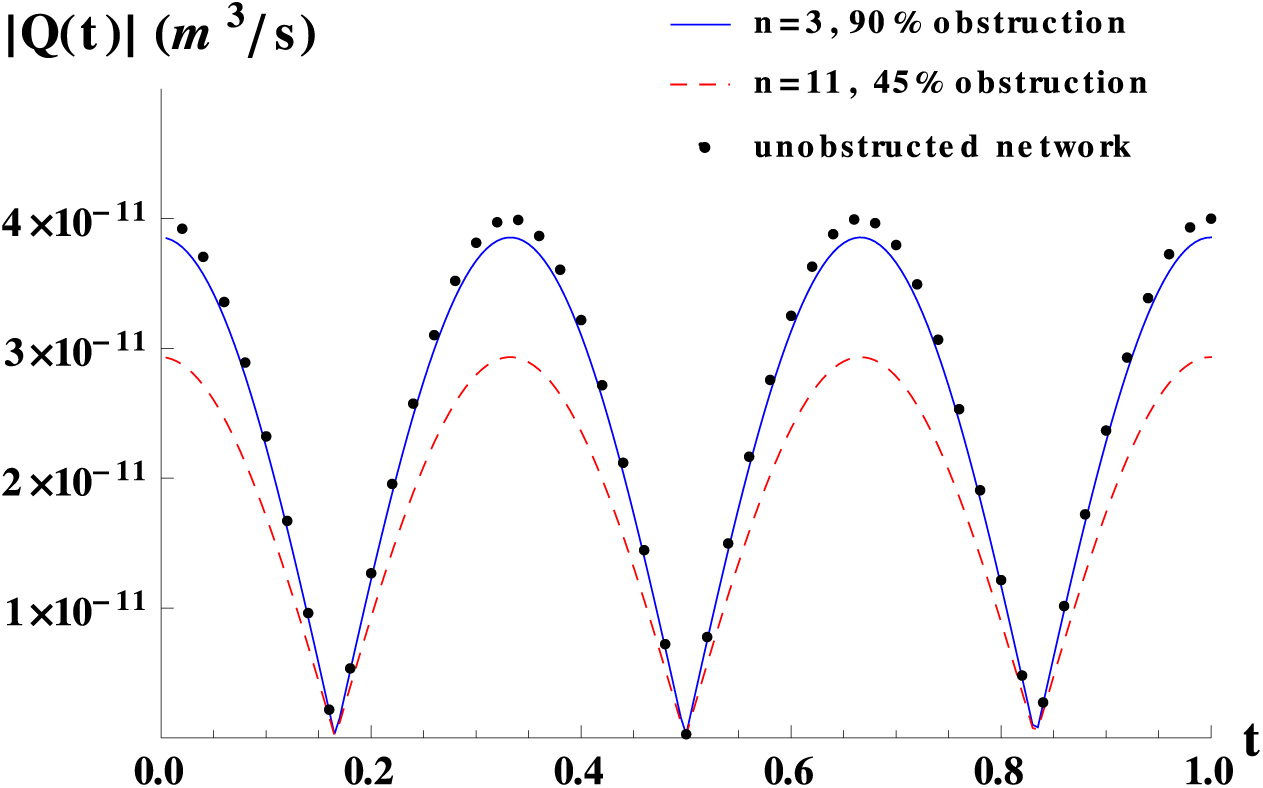
Time-dependent flow for obstructed networks with a jump in resistance. Time-dependent flow for a network obstructed by 90% in area at level 3, for a network obstructed by 45% in area at level 11 and for a network without obstructions as reference. The network used was the same as in Fig. 8. The total pressure drop was set to 600 Pa.

Despite the tendency shown in Fig. 8A that seems to imply that the larger the obstruction, the larger the jump Δ*χ*, this does not hold for very large degrees of obstruction, i.e. when *f ≈* 1, Δ*χ ≈* 0, a regime that may be physiologically relevant only for vessel suppression. As illustrated in Fig. 8B, in this regime the network response changes dramatically with respect to its unobstructed network counterpart, independently of the position of the obstruction relative to the resistance jump. It is worth noticing that, as depicted in Fig. 8A, a minimum in the response can be observed when occlusions take place at the same position that the resistance jump.

## Conclusions

In this work we relate network structure to hemodynamics after occlusions. We study tree-like networks and find that when vessels are equal, the drop in the dynamic response, after obstructions on half of its vessels at a given bifurcation level, increases monotonically with the bifurcation level in which they occur. The outer the level where the obstructions are included, the larger the drop in the dynamical response of the network.

For vessels that follow Murray’s law for the sequence of radii, and a power law for the sequence of lengths, two radically different behaviors are observed for the response function: one for large arteries, in which the response increases as a function of the level *n* in which obstructions are located, and one for smaller vessels, in which the response decreases as a function of the level *n* in which obstructions are placed. This example shows the non-trivial effect of vascular obstructions and puts forward the importance of analytic equations in the interpretation of the numerical results.

For networks in which jumps in resistance between subsequent levels exist, we have identified the sites of the jumps as critical for the overall capacity of the network to supply blood to a tissue. We have also demonstrated that relatively small obstructions in these critical topological sites, cause a much larger decrease on flow than larger obstructions in non-critical sites. By simple observation of the structure of a vascular network, these key sites could be readily identified and monitored in vivo.

We have derived analytical expressions for the dynamic response of the network that provide insight into the relevant mechanisms that control flow across a tree-like network in the presence of vessel occlusions. Moreover, we are able to find approximations for cases that might be physiologically relevant in which the roles played by the network structure, the degree of obstruction and the geometrical place in which obstructions take place can be clearly identified. We demonstrate that our results are local in the sense that they depend on the network structure around the place where obstructions are. This implies that whenever there is a tree-like network in an in-vivo vasculature, our model is able to interpret the effect that an obstruction has on flow.

Our results demonstrate that the effect of occlusions on flow is somehow opposite to the effect of the addition of anastomotic vessels [14]. When in the former case occlusions cause a decrease on flow, in the latter, anastomosis causes an increase on flow. However they are similar in the sense that whenever vascular alterations to a network occur (either in the form of occlusions or in the form of addition of vessels), the impact on flow has similar tendencies. Namely, for vascular networks with *a <* 2 (*a >* 2), the outer (inner) the vascular alterations are placed, the larger the impact on flow. Also, sites presenting a large jump in resistance are critical for the overall capacity of the network to supply blood. In the case of obstructions situated on those critical sites, a large decrease in network response (and flow) might compromise the survival of the tissue. While new vessels in the form of anastomosis added (naturally or artificially) at those critical sites, result in an extremely large increase in the network response and flow.

For our model, we have considered that vessels are rigid. However, real vessels are elastic tubes. Important insight could be gained from the inclusion of elastic effects in the model, especially for large arteries, where elastic effects are very important. Regional tissue metabolism, such as the myogenic effect in arteries, can be considered, as was done in reference [14], where it was shown that the impact that is has on flow is large, but tendencies with the geometrical place where anatomical variations occur, are qualitatively unaltered.

A validation of our model with an in-vivo biological system is currently not possible, since it would require data of flow measurements after a systematic variation of the anatomical place where obstructions are located. Such information does not exist in literature. One possibility to validate the model, at least at certain length scales, in an experimental situation, would be to use artificial vessel networks, such as networks etched in microfluidic devices, where local variations could be systematically done.

Vascular alterations might affect or help a patient depending on the medical condition, in the case of ischemic conditions affecting the irrigation of vital tissues, the presence of occlusions is negative and the presence of anastomosis is positive for tissue irrigation. On the contrary, for a vasculature irrigating a tumor, anastomosis might be good for the tumor and bad for the patient and occlusions would have the opposite effect. Our results can help to decide the anatomical sites where selective suppression of vessels, might help to decrease blood flow towards the tumor, decreasing its oxygen supply. The study of the effect on flow of other vasculature alterations, such as the presence of aneurisms, is desirable.

Our results put forward the importance of relating vasculature structure with hemodynamics. Recent progresses on high resolution microscopy allows the visualization of the characteristics of an individual vasculature [34]. Hence, the concurrence of sophisticated image-tracking systems and mathematical models such as the one presented here, provides a new perspective and relevant tools in the determination of blood supply to a tissue.

**Supporting Information**

**S1 Supporting Information**

**Supplementary material**

## References

1. Schwartz CJ, Gerrity RG. Anatomical pathology of sudden unexpected cardiac death. Circulation. 1975;52(6 Suppl): III18–26.

2. Hayreh SS, Kolder HE, Weingeist TA. Central retinal artery occlusion and retinal tolerance time. Ophthalmology. 1980;87: 75–78.

3. Hayreh SS, Jonas JB. Optic disk and retinal nerve fiber layer damage after transient central retinal artery occlusion: an experimental study in rhesus monkeys. Am J Ophthalmol. 2000;129: 786–795.

4. Stergiopulos N, Young DF, Rogge TR. Computer simulation of arterial flow with applications to arterial and aortic stenoses. J Biomech. 1992;25: 1477–1488.

5. Stroud JS, Berger SA, Saloner D. Influence of stenosis morphology on flow through severely stenotic vessels: implications for plaque rupture. J Biomech. 2000;33: 443–455.

6. Botnar R, Rappitsch G, Scheidegger MB, Liepsch D, Perktold K, Boesiger P. Hemodynamics in the carotid artery bifurcation: a comparison between numerical simulations and in vitro MRI measurements. J Biomech. 2000;33: 137–144.

7. Long Q, Xu XY, Ramnarine KV, Hoskins P. Numerical investigation of physiologically realistic pulsatile flow through arterial stenosis. J Biomech. 2001;34: 1229–1242.

8. Formaggia L, Lamponi D, Tuveri M, Veneziani A. Numerical modeling of 1D arterial networks coupled with a lumped parameters description of the heart. Comput Methods Biomech Biomed Engin. 2006;9: 273–288.

9. Varghese SS, Frankel SH, Fischer PF. Direct numerical simulation of stenotic flows. Part 1. Steady flow. J Fluid Mech. 2007;582: 253–280.

10. Varghese SS, Frankel SH, Fischer PF. Direct numerical simulation of stenotic flows. Part 2. Pulsatile flow. J Fluid Mech. 2007;582: 281–318.

11. Frauenfelder T, Boutsianis E, Schertler T, Husmann L, Leschka S, Poulikakos D, et al. In-vivo flow simulation in coronary arteries based on computed tomography datasets: feasibility and initial results. Eur Radiol. 2007;17: 1291–1300.

12. Lee SE, Lee SW, Fischer PF, Bassiouny HS, Loth F. Direct numerical simulation of transitional flow in a stenosed carotid bifurcation. J Biomech. 2008;41: 2551–2561.

13. Alastruey J, Parker KH, Peiró J, Byrd SM, Sherwin SJ. Modelling the circle of Willis to assess the effects of anatomical variations and occlusions on cerebral flows. J Biomech. 2007;40: 1794–1805.

14. Flores J, Meza Romero A, Travasso RDM, Corvera Poiré E. Flow and anastomosis in vascular networks. J Theor Biol. 2013;317: 257–270.

15. Suda M, Eder OJ, Kunsch B, Magometschnigg D, Magometschnigg H. Preoperative assessment and prediction of postoperative results in an artificial arterial network using computer simulation. Comput Methods Programs Biomed. 1993;41: 77–87.

16. Grasman J, Brascamp JW, Van Leeuwen JL, Van Putten B. The Multifractal Structure of Arterial Trees. J Theor Biol. 2003;220: 75–82.

17. Flores J, Corvera Poiré E, del Río JA, López de Haro M. A plausible explanation for heart rates in mammals. J Theor Biol. 2010;265: 599–603.

18. Thurston GB, Henderson NM. Effects of flow geometry on blood viscoelasticity. Biorheology. 2006;43: 729–746.

19. Morrison FA. Understanding Rheology. New York: Oxford University Press; 2001.

20. Bravo-Gutiérrez ME, Castro M, Hernández-Machado A, Corvera Poiré E. Controlling viscoelastic flow in microchannels with slip. Langmuir. 2011;27: 2075–2079.

21. Collepardo-Guevara R, Corvera Poiré E. Controlling viscoelastic flow by tuning frequency during occlusions. Phys Rev E Stat Nonlin Soft Matter Phys. 2007;76: 026301-1–026301-7.

22. Nichols WW, O’Rourke MF. McDonald’s Blood Flow in Arteries Theoretical, Experimental and Clinical Principles. New York: Arnold/Oxford University Press; 1998.

23. The low frequency results are indistinguishable from steady-state results. However, we consider important to keep the formalism as general as possible in order to make it applicable when external frequencies are applied to a network [21].

24. Kern TS, Engerman RL, Peterson CM. Elevated blood viscosity in alloxan diabetic dogs and experimentally galactosemic dogs. J Diabetes Complicat. 1989;3(3): 158–162.

25. Experimental relaxation times for human blood, might vary several orders of magnitude, depending on confinement and dynamic conditions [18]. For the network studies presented in this paper, the dynamic response is almost independent of the choice of this quantity since we work at low frequencies.

26. It is worth pointing out that at level *i* + 1, there are twice the number of vessels than at level *i* and the flow is halved.

27. Murray CD. The physiological principle of minimum work. I. The vascular system and the cost of blood volume. Proc Natl Acad Sci USA. 1926;12: 207–214.

28. Sherman TF. On connecting large vessels to small. The meaning of Murray’s law. J Gen Physiol. 1981;78: 431–453.

29. Williams HR, Trask RS, Weaver PM, Bond IP. Minimum mass vascular networks in multifunctional materials. J R Soc Interface. 2008;5: 55–65.

30. Taber LA, Ng S, Quesnel AM, Whatman J, Carmen CJ. Investigating Murray’s law in the chick embryo. J Biomech. 2001;34: 121–124.

31. McCulloh KA, Sperry JS, Adler FR. Water transport in plants obeys Murray’s law. Nature. 2003;421: 939–942.

32. McCulloh KA, Sperry JS, Adler FR. Murray’s law and the hydraulic vs mechanical functioning of wood. Funct Ecol. 2004;18: 931–938.

33. McCulloh KA, Sperry JS. The evaluation of Murray’s law *In Psilotum Nudum* (Psilotaceae) an analogue of ancestral vascular plants. Am J Bot. 2005;92(6): 985–989.

34. Zhang L, Hu C, Zhao T, Luo S. Noninvasive visualization of microvessels using diffraction enhanced imaging. Eur J Radiol. 2011;80: 158–162.

